# Effect of ring topology in a stochastic model for Z-ring dynamics in bacteria

**DOI:** 10.1101/452441

**Authors:** A. Swain Sumedha, A. V. A Kumar

## Abstract

Understanding the mechanisms responsible for dynamics of the *Z*-ring is important for our understanding of cell division in prokaryotic cells. In this work, we present a minimal stochastic model that qualititatively reproduces observations of polymerization, of formation of dynamic contractile ring that is stable for a long time and of depolymerization shown by FtsZ polymer. We explore different mechanisms for ring breaking and hydrolysis. Hydrolysis is known to regulate the dynamics of other tubulin polymers like microtubules. We find that the presence of the ring allows for an additional mechanism for regulating the dynamics of FtsZ polymers. Ring breaking dynamics in the presence of hydrolysis naturally induce rescue and catastrophe events, irrespective of the mechanism of hydrolysis. Based on our model, we conclude that the *Z*-ring undergoes random breaking and closing during the process of cell division.

## 1 Introduction

Cell division is one of the most fundamental processes in living cells. In prokaryotic cells numerous proteins take part in assembling the machinery of cell division, called the divisome. FtsZ(Filamenting temperature-sensitive mutant Z) is the most important among them. This tubulin homologue polymerizes head to tail, forming dynamic protofilaments. These protofilaments together form the core of a structure called the Z-ring in the division plane of the cell[1-5]. The Z-ring persists throughout the cell division. Recent studies on septal wall suggest that FtsZ monomers move around the ring via treadmilling which guides and regulates their growth, which in turn controls division [6-10]). The dynamic ring is a phenomena that is intrinsic to FtsZ and the importance of this contractile ring in cell division is well established by now. Recently, there have been a lot of focus both invivo[8, 9] and invitro[11, 12] to understand the dynamics of the ring. But the mechanism controlling the activity of the ring is poorly understood.

The monomers of FtsZ consists of two independent domains. The *N*-terminal domain with its parallel beta sheets connected to alpha helices, provides the binding site for GTP/GDP. The C-terminal region is essential for FtsZ to interact with other proteins like FtsA and ZipA[13] and also as a membrane tether for the Z-ring to form during cell division[14]. Above a critical concentration of GTP, FtsZ polymerizes cooperatively into filaments which are single stranded and have a head to tail orientation with polymerization at only one end [15-24]. The exact mechanism behind this cooperativity is not well understood. Studies have shown that FtsZ can polymerize into multi-stranded bundles, sheets and form chiral vortex on membranes [10, 24–29] depending on the assembly buffer, nucleotide and other proteins present.

Atomic Force Microscopy(AFM) of filaments adsorbed on Mica surface have shown the role played by lateral interactions and filament curvature in the understanding dynamic behavior of FtsZ filaments[30, 31]. The study of individual filaments under AFM showed that the filaments can form rings which are dynamic and may open up, lose material and close again until they open irreversibly and then the filaments depolymerize. It has been shown that unlike the case of tubulin, where nucleotide exchange is the rate limiting step for the reaction kinetics, in the case of FtsZ it is the hydrolysis which controls the reaction kinetics. The ring formation of FtsZ is studied in presence of GTP, guanosine-5-[(β, γ)-methyleno-triphosphate (GMPCPP) which reduces the hydrolysis as well as depolymerization rates and GTP with glycerol in the buffer which reduces the depolymerization rates and hence stabilize the lifetimes for irreversible opening[12]. They found that the rings formed in the presence of GMPCPP had broader length distribution, with higher average length than the rings formed in the presence of GTP. A very recent study by Diego et al [11] has again focused on the ring formation invitro. They made some key observations. They found that the fine tuning of hydrolysis is needed for the formation of stable rings. They also found that the treadmilling results from a directional growth of curved and polar filaments from the nucleation point at the membrane. The preferential addition of GTP subunit to the leading edge establishes a GTP-GDP gradient.

FtsZ monomers are very similar to eucaryotic tubulin polymers like microtubules(MTs) and actin in their composition and functioning. MTs are semiflexible polymers that are a key component of the mitotic spindle[32] in eukaryotic cells and this requires them to be dynamic in nature. They exhibit a phenomena of dynamic instability(DI), in which they switch from a phase of slow elongation to rapid shortening (catastrophe) and from rapid shortening to growth(rescue). A polymerizing MT grows until it suffers a catastrophe and starts to depolymerise. Similarly, a depolymerising MT undergoes rescue and starts polymerising again. Attempts using stochastic models have been successful in explaining this behaviour[33–37]. Experimental results and theory together have now established that the hydrolysis of tubulin monomers is responsible for the dynamic instability of MTs. Though the exact mechanism of hydrolysis is still debated [38]. Similarly, actin filaments[39, 40] also exhibit non equilibrium phenomena known as treadmilling. In treadmilling, the new subunits get added at the growing end and old subunits leave the polymer from the other end. Mathematical models of MTs and actin have helped our understanding of the phenomena of dynamic instability and treadmilling. These models bridge scales and hence facilitate our understanding of complex biological phenomena in terms of elementary processes [41]. This helps in organizing the plethora of information one gets from the biochemistry study of these proteins.

The basic self-assembly mechanism underlying DI, assembly mediated by nucleotide phosphate activity, is omnipresent in biological systems. FtsZ falls in the same class of bio-polymers and hence it seems natural that a similar modelling approach capturing the dynamics will help our understanding of the FtsZ. But except for a few deterministic approaches to model the FtsZ ring, there have been almost no attemps at modelling FtsZ at the molecular level. Most of these deterministic models aimed to characterise thoeretically the in vivo and in vitro observations of FtsZ assembly. These models involve a number of differential equations to be solved simultaneously making it computationally very costly. The eight equation model proposed by Chen *et al*. [42, 43] described the initial stages of FtsZ polymerization successfully, but fails to handle the whole process of FtsZ assembly. There are other models by Dow *et al*. [44], Lan *et al*. [45] and Surovtsev *et al*. [46] which employ few hundreds of differential equations making the computations very complex. Recently, kinetic models based on average charcteristics of different species and their concentrations have been proposed, where the number of differential equations need to be solved has reduced considerably to 17[47] or 10[48]. These models reduced the computational cost drastically and were able to predict the time taken to reach the steady state, the concentration of FtsZ in the Z-ring and average dimension of the filaments and bundles, which were in agreement with the experimental observations. However, the dependence of these on factors like rate of hydrolysis could not be obtained by these models. These deterministic models cannot capture the ring dynamics, which is stochastic in nature.

MTs and actin form straight filaments while FtsZ form a ring of roughly one micro-meter diameter in its active state [11]. In this paper we propose a stochastic model for FtsZ which is similar in spirit to known models of MTs and actin with new additional processes to account for ring formation and breaking. We consider both possible mechanisms of hydrolysis of tubulin monomers: namely the vectorial and stochastic hydrolysis [38]. In the stochastic hydrolysis, GTP-FtsZ subunit can hydrolyze in a stochastic manner, irrespective of the position of the subunit in the protofilament. The rate of hydrolysis in this case would be proportional to the amount of unhydrolyzed monomers. In vectorial hydrolysis, an unhydrolyzed monomer gets hydrolyzed only if the neighboring monomer is already hydrolyzed. This is a highly cooperative mechanism and there exists a sharp boundary between the hydrolyzed and unhydrolyzed parts of the FtsZ filaments. We find that the hydrolysis of the monomers is essential for the dynamics of FtsZ polymer. But interestingly the rescue and catastrophe events are more crucially regulated by the process responsible for opening and closing of the Z-ring. In this paper, we introduce a model which takes into account both the processes: hydrolysis and ring breaking. We find that the random breaking of ring naturally makes the FtsZ polymer more dynamic. The stochastic model outlined in this article is much simpler and computationally less costly. Moreover it captures the essential features of FtsZ dynamics observed in the experiments and provides fresh inputs to the understanding.

When polymerized FtsZ in a solution of GTP is exposed to an excess of GDP it depolymerizes quickly[49]. GTP-bound monomers have been found to constitute 80% of filaments under in vitro conditions[17, 50]. In our model we assume that there is no polymerization of GDP-bound monomers. Though GDP-bound monomers have been found to polymerize in vitro but as compared to polymerization of GTP-bound monomers the equilibrium constant for this process has been found to be significantly lower[20]. The GDP-bound monomers are also found to be relatively shorter and more curved than their GTP-bound counterparts[51]. These results suggest that the GDP-bound polymerization is unlikely to be viable and thus irrelevant in explaining FtsZ dynamics.

Our stochastic model incorporates the processes happening chemically at a monomer level and try to explain in vitro observations for isolated filaments. In our study, we consider both hydrolysis mechanisms, namely: vectorial hydrolysis(VH) and stochastic hydrolysis(SH) and find the conclusions largely insensitive to the nature of hydrolysis. We also consider two different processes for ring breaking:a) rupture can occur at any monomer-monomer interfaces. This we call random breaking,b) rupture can occur only at the interface with atleast one GDP bound FtsZ monomer. We call this non-random ring breaking. We consider four scenarios in this paper based on the nature of hydrolysis and the mechanism of ring breaking: vectorial hydrolysis with random ring breaking, vectorial hydrolysis with non-random ring breaking, stochastic hydrolysis with random ring breaking and stochastic hydrolysis with non-random ring breaking. The stability of ring improves with randomness and stochastic hydrolysis with random ring breaking gives the most stable rings. But we find that the experimental observations can be successfully explained by the vectorial hydrolysis with random ring breaking and hence the non-random ring breaking mechanism can be ruled out. This agrees with earlier suggestions [12] and also with the suggestions that ring breaks due to tension created on the ring due to deformation of the membrane [57, 58]. Ours is the first microscopic stochastic model for the study the *Z*-ring dynamics.

The plan of the paper is as follows: In Section 2 we describe our model. In Section 3.1 we consider the vectorial hydrolysis with random and nonrandom breaking of the ring and find that random ring breaking gives rise to stable dynamic ring. In section 3.2 we study the effect of changing the ring breaking rate on the dynamics and conclude that with random ring breaking one gets a contractile ring. In section 3.3 we look at the stochastic hydrolysis with random and non-random ring breaking and compare it with the results of vectorial hydrolysis. We discuss our results in Section 4.

## 2 Methods

### 2.1 The Model

FtsZ filament is known to exist both in the form of an open chain and as a closed ring. In the open form it has a single active end(the arrow head) which is in contact with a reservoir with GTP-bound monomers. It is considered to be directional with polymerization only possible at its head end. In our model, in the presence of a GTP-bound monomer at its polymerizing end, a GTP-bound monomer gets added to the polymer at a rate λ. Whenever a GDP-bound monomer is at either ends of an open filament, the polymer can lose a GDP-bound monomer from the ends with a rate *μ*. Both these processes are independent. It has been experimentally observed that depolymerization is faster at the ends as compared to the center[12]. Hence, we have assumed depolymerization only at the ends. Also, in general the polymerisation rate is higher than the depolymerization rate[10].

Our minimal stochastic model incorporates the process of polymerization at the preferred end, depolymerization from both ends, hydrolysis, ring breaking and closing. In Figure 1, we present the schematic of all the processes we include in our model.

**Figure 1.**
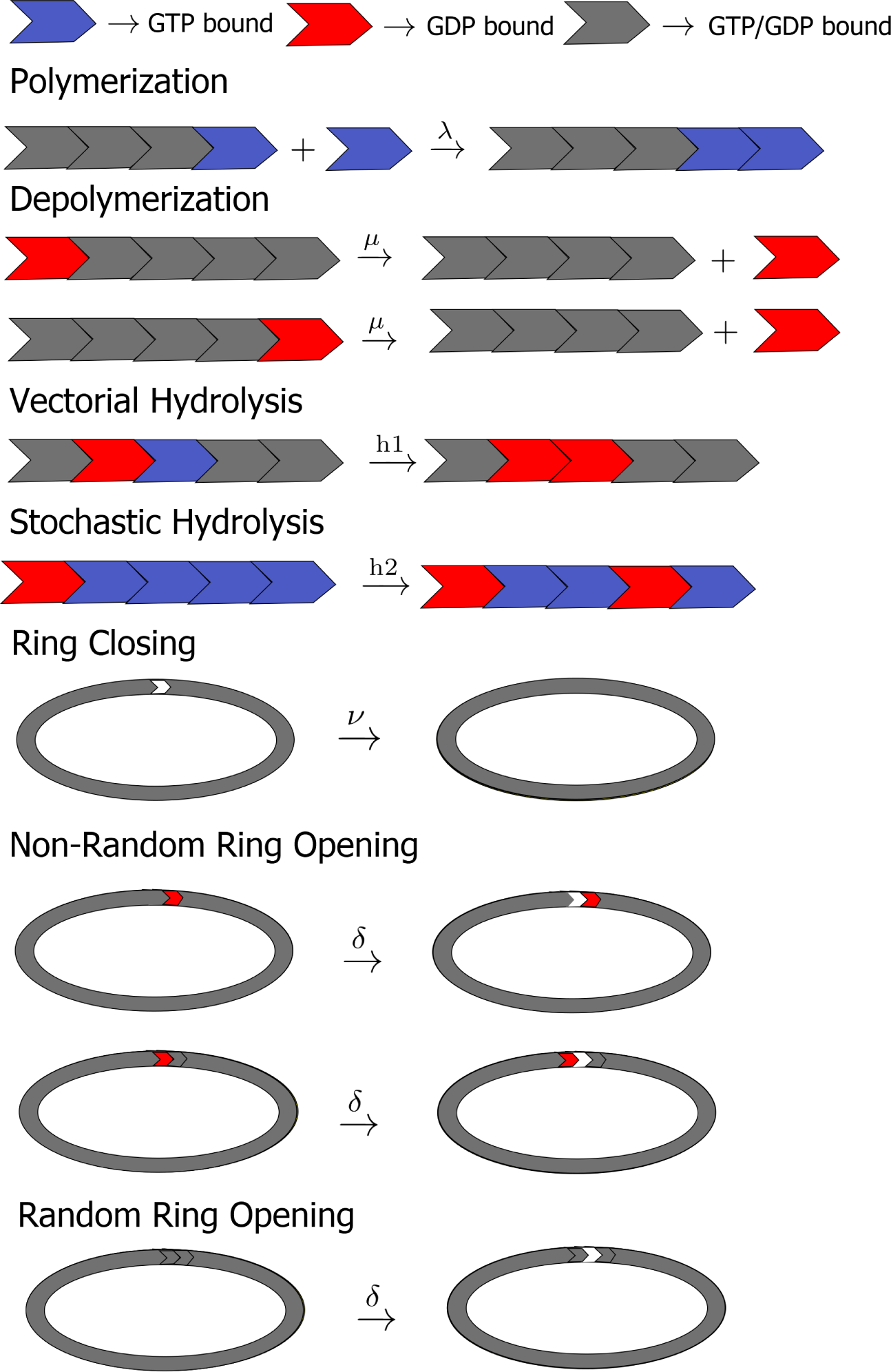
Schematics for the different reactions taking place in the system.

The cases for both vectorial and stochastic hydrolysis have been considered individually. In the case of vectorial hydrolysis in the presence of a GDP-GTP interface, the interface grows by converting the GTP-bound monomer associated with it to a GDP-bound monomer with a rate *h*_1_. Propensity of a reaction in stochastic reactions gives us the likelihood of a particular reaction happening in a unit time. Reactions with higher propensities are more likely. Thus the propensity for this reaction is

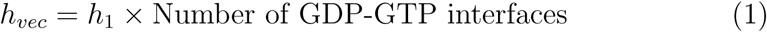

In the case of stochastic hydrolysis, a GTP-bound monomer can get randomly converted into a GDP-bound monomer with a rate *h*_2_. The propensity for the reaction is

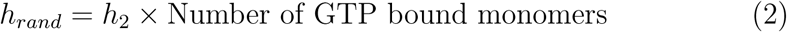

The hydrolysis reactions can take place both in the open as well as ring forming filaments.

Following the scheme used in earlier works[46, 52] to model the natural curvature of FtsZ we assume that the filament has an optimum length(*N*_0_) at which it has the maximum probability of closing and forming a ring. Hence, given a length *N*, the polymer has a non-zero probability to form a ring. We assume the rate of this process depends on *N* and is given by a Gaussian distribution that peaks at *N* = *N*_0_:

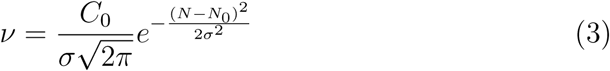

Ring opening was assumed to have two possibilities. One in which the ring opening is completely random. If we take δ to be the rate of random breaking, then *U*_2_ = *δN* is the propensity of this reaction. Given that the GDP associated bonds are much weaker than GTP associated bonds, we also consider the case where the ring breaking is only possible at an interface containing atleast one GDP-bound monomer. So, for a given ring with *N*_1_ GDP bound interfaces, the propensity of breaking is *U*_3_ = *δN*_1_.

### 2.2 Algorithm

The dynamics hence can be described by a set of coupled chemical master equations for the probability of open and closed polymer. Due to the presence of ring breaking, it is not possible to solve the equations even in the simpler case of vectorial hydrolysis. Hence we use Gillespie algorithm[53, 54] to solve the chemical master equations numerically. Gillespie algorithm offers an elegant way to speed up simulations by doing away with the many rejected trials of the traditional Monte-Carlo moves. While, traditional Monte-Carlo methods check at each step if each reaction takes place, Gillespie algorithm draws directly the time elapsed until the next reaction or process occurs and which reaction takes place at that time. The advantage of Gillepsie algorithm is that it generates an ensemble of trajectories with the correct statistics. It has been very successful in simulating many chemical and biological reactions [54].

## 3 Results and Discussion

### 3.1 Vectorial Hydrolysis

In this section we study the dynamics when the hydrolysis is assumed to be only at the interface, i.e the vectorial hydrolysis. Let *P_r_* (*N*, *t*) represent the probability of having a closed ring polymer of length *N* at time *t* and *P_o_*(*N*, *t*) represent the probability of having an open chain polymer of length *N* at time *t*. We define *p*_1_(*t*) as the probability of having a GTP-bound monomer at the active end in the open ring configuration. Thus the probability of active end having a GDP bound monomer is 1 − *p*_1_(*t*). Let *p*_2_(*t*) be the probability of finding a GDP bound monomer at the non-polymerizing end. The probability of observing a GDP monomer at either end is *p*_3_(*t*) = *p*_2_(*t*) + 1 − *p*_1_(*t*), Also as discuseed in Section 2, 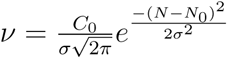 is the rate for an open polymer of length *N* to close up into a ring. Then for random breaking the system can be described by the following set of equations:

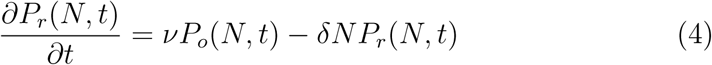

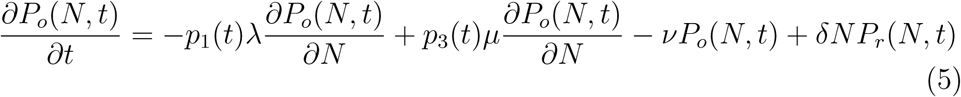

In the beginning there are only the processes of polymerization, depolymerization from the back end and hydrolysis. Once the polymer takes a ring form, it doesn’t have a polymerizing and a non-polymerizing end. In this state for sometime there are only two reactions going on:hydrolysis and ring opening. Thus polymerization rate does not have a direct effect on the dynamics of a closed ring. The ring opening being a random process, it is not possible to tell what will the ends of the polymer be when it opens up. Once the ring opens up, it can again undergo polymerization and depolymerization. Depending on the tip of the open polymer, the polymer undergoes dynamic catastrophe and rescue events. This makes the ring length and the ratio of GDP/GTP bound monomers in the ring a fluctuating parameter, which makes calculating *p*_1_(*t*) and *p*_3_(*t*) impossible. Hence, it is not possible to solve the chemical master equations exactly. One can solve the equations exactly only in the trivial case where hydrolysis is much lower than the polymerization rate and hence the probability *p*_1_(*t*) and *p*_3_(*t*) can be taken to be 1 and 0 respectively. In this case one gets a gaussian distribution for the ring length distribution. We hence simulate the system using Gillespie algorithm described in Section 2. We will compare the mechanisms of ring opening i.e random breaking and non-random breaking by comparing the properties of experimentally possible observables like ring length and ring life time. We fix depolymerisation rate to be 1 (as we can always fix one of the rates to be one) and take *N*_0_ = 120 and *σ* = 15. *C*_0_ is chosen such that 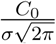 is 10. We will now study the effect of polymerisation,hydrolysis and ring breaking rate on the dynamics of the ring.

In figure 2(left) we plot length of FtsZ polymer as a function of time from a typical run for different hydrolysis rates for random breaking of FtsZ rings. We fix the random breaking rate to be *δ* = 0.008. From the figure, it is clear that there are three distinct regions. Initially, there is a growth phase, where the polymer chain is formed from FtsZ monomers. In this phase, the length of the polymer grows almost linearly with time. This phase is insensitive to hydrolysis rate, *h*_1_. After the initial growth phase, the polymer makes a transition to a phase where the length gets stabilised for a period. This is the phase where FtsZ ring exists. It is clear from the trajectories that FtsZ ring is dynamic as the length fluctuates with time. The third region is where the polymer either grows an open chain or depolymerise completely.

**Figure 2.**
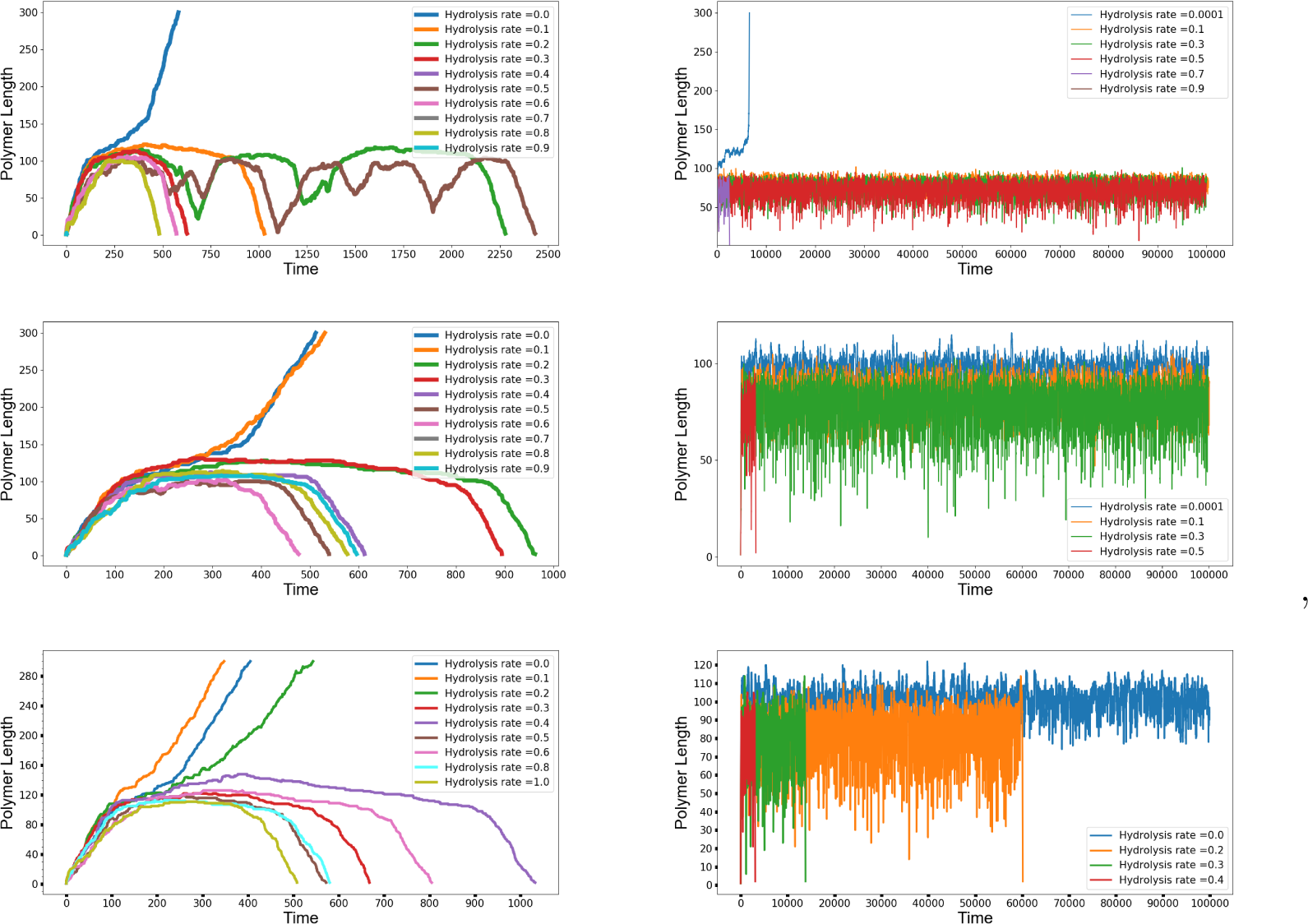
Trajectories for Random(left) and Non-Random Breaking(right) for polymerization rate 2.0, 4.0 and 8.0 respectively(top to bottom)

When we consider random breaking, ring can break at any of the interface. Since *h*_1_ = 0, there is no hydrolysis taking place, there are no GDP bound monomers in the polymer. So when the ring opens up, it cannot depolymerise. Hence, after a short period of dynamic ring formation and breaking, the chain grows as a linear chain. When the hydrolysis rate is not zero, but small there is finite probability to encounter a GDP bound monomer at either of the ends whenever the ring opens up. In this case, depolymerisation occurs before the chain start polymerising again resulting in a larger lifetime for the dynamic ring. As the hydrolysis rate increases, the probabilty of finding GDP bound monomer increases. In the case where GDP bound monomers are present at the polymerizing end, all of them need to get depolymerized to expose a GTP bound monomer for the polymer to grow again. We find that this results in rescue and catastrophe events in which the polymer may lose as many as 20-30 monomers before recovering again. Additionally, the opening and closing dynamics ensures that we have a lot many interfaces in this case. Finally, when the polymer has no more GTP bound monomers and only has GDP bound monomers there are no more rescue events and the polymer depolymerizes completely. Thus at very low hydrolysis rates the polymer forms a ring for a short time and then escapes. We find that as hydrolysis rate increases, initially the ring life time increases. Because of interplay between polymerization, depolymerization, ring closing and opening, ring life time peaks at a finite value of *h*_1_ and beyond that stability of ring keeps on decreasing.

In Figure 2(right), we plot the length of FtsZ polymers in the case of nonrandom breaking; i.e. the ring opens only at the interface formed by atleast one GDP bound monomer, for different hydrolysis rates. In the zero hydrolysis case once the ring gets formed it will never open as there is no GDP bound interface present. At small values of hydrolysis rate, GTP will get hydrolyzed after a long time and at some subsequent time it will open. Since GDP bound monomers are present at least at one of the ends, the chain is exposed for depolymerisation once it opens up. Hence, chances of depolymerization is bigger in this case in contrast to the random breaking case. This results in the large fluctations seen in the trajectories compared to the case of random breaking. The number of interfaces for vectorial hydrolysis remains the same, i.e, 1 throughout the time evolution(recall that for random breaking the number of interfaces increase with time). In some rare cases for very high polymerization rates the ring does escape to form a open chain.

From figure 2, it is clear that the ring dynamics and stability is sensitive to the hydrolysis rate in the case of random breaking of the ring and insensitive to hydrolysis rate in the case of non-random breaking. These observations can be made more quantitative in terms of experimentally measurable quantities like ring lifetime and length distributions. We obtain these quatities by averaging over 10^5^ independent trajectories.

#### 3.1.1 Ring length distribution

In Fig 3(a) and 3(b) we plot distribution of ring size in both random breaking and non-random breaking respectively for different hydrolysis rates. In the case of random breaking the distribution gets narrower with increasing hydrolysis rate. In ref. [12] authors studied the ring length distribution of FtsZ polymers in-vitro. They found that rings formed in the presence of slowly hydrolyzing analogue, GMPCPP, resulted in broader distribution, which was shifted to the right. Our plots in Fig. 3(a) are consistent with this observation. The ring length distribution gets thinner with increasing hydrolysis rate and the peak shifts to the left, resulting in decreasing average length of the ring. In contrast, in the case of the non-random ring breaking, except for zero hydrolysis rate, the ring length distribution remains unchanged, suggesting that the ring length is insensitive to the changes in hydrolysis rate. In the case of zero hydrolysis rate, the ring never opens up once it forms as there are no GTP-GDP interfaces, and the distribution is just proportional to the ring closing rate *v*.

**Figure 3.**
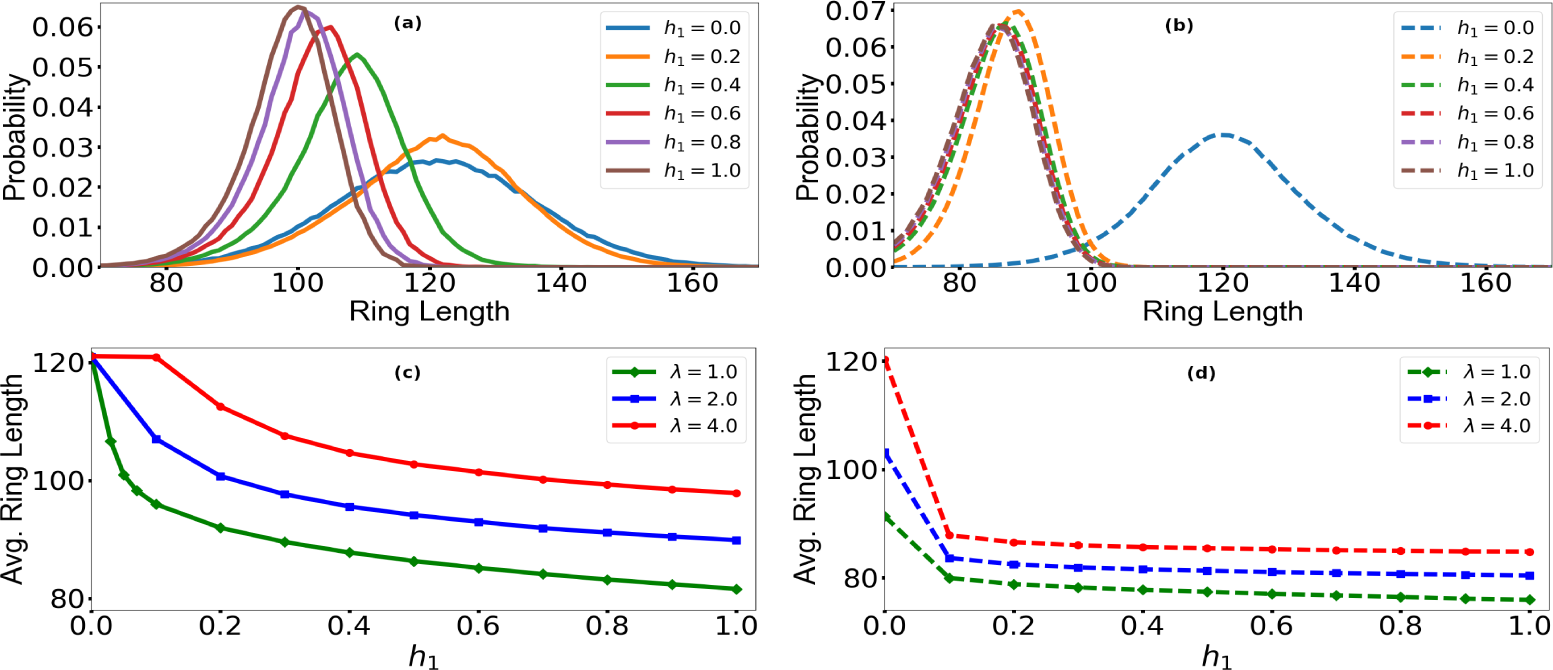
a) and b): Plot of distribution of ring length for polymerization rate 4.0 for random and non-random ring breaking respectively. c) and d): Plot of average length for different polymerization rates, for random and non-random breaking respectively

In fig 3(c) and 3(d) we plot the average length in the two cases. As the hydrolysis rates increased, the average length of the polymer was found to decrease gradually in the case of random breaking. In the case of nonrandom breaking, the average length remains same except for the case of zero hydrolysis rate. This suggests that the observation that the dynamics of FtsZ polymer is insensitive to the hydrolysis rate in the case of non-random breaking, made based on figure 2(b) is indeed true. Near zero hydrolysis rate, the average length of ring, depends only on the polymerisation rate as the number of GDP-bound monomers are very less and depolymerisation does not take place often. So the average length of the ring increases as the rate of polymerization increases.

#### 3.1.2 Time for irreversible ring opening

It is known that the dynamic ring stage of FtsZ polymer is very stable. The ring undergoes many catastrophe and rescue event during this time, before eventually decaying. Hence, one of the most important observable is the total time for which the ring exists before it eventually escapes the ring formation and grows as a linear chain or decays completely. Average time for this irreversible ring opening has been plotted in Figures 4(a) and 4(b) for random and non-random ring breaking respectively. For random breaking it peaks at a finite value of hydrolysis rate. This is because at very low values of hydrolysis the polymer escapes into growth phase and at large value of hydrolysis, the chances of ring closing again is small. Hence there is a narrow window of hydrolysis rate where the dynamic ring is most stable. This is consistent with the very recent experiments of Diego et al [11] where they found that fine tuning of hydrolysis was needed to get stable dynamic rings. Moreover, we observe that the region of maximum stability occurs roughly at a fixed ratio of hydrolysis to polymerization rates. It has been observed that in the presence of a GTP regeneration system, the dynamic ring can continue for a long time[49]. This along with the fact that FtsZ remains highly conserved across species[55], can be used to argue that FtsZ sits on the edge of stability in its naturally occurring form. This is clearly demonstrated in the plots for random ring opening in which, across all polymerization rates the region of maximum stability happens at a fixed ratio of hydrolysis and polymerization rates. In contrast to the case of random ring opening, for non-random breaking the ring is not really dynamic and hence the lifetime goes down monotonically with the hydrolysis rate.

**Figure 4.**
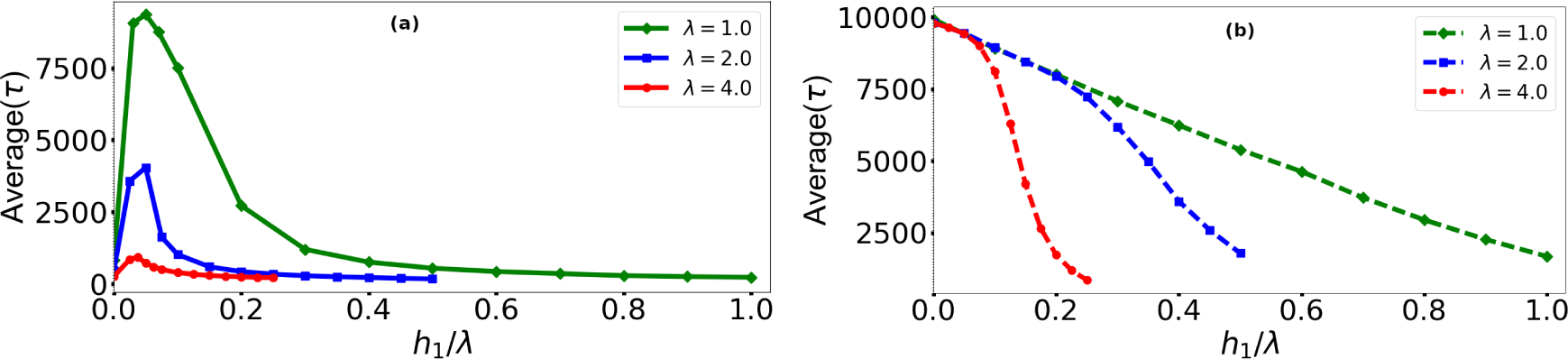
Average lifetime(τ) for irreversible opening for random(a) and non-random(b) breaking. Here τ is the time for irreversible opening.

#### 3.1.3 Ring lifetime distribution

We can also look at the distribution of time elapsed between two openings. We expect that there would be many small events where the ring opens with GTP at the growing end and close back immediately. Figure 5(a) and 5(b) shows these distributions for the case of random and non-random ring breaking. The distributions become flatter and flatter as the hydrolysis rate increases for both cases; however the effect is more pronounced in the case of non-random breaking. Polymerization rate does not affect the lifetime of single openings since the polymerisation does not take place when the ring is closed. Such a trend can be observed in the case of random ring opening. But in case of non-random opening, increase in polymerization rate leads to an decrease in average ring lifetime for the same hydrolysis rate. Hence, we conclude that in the case of non-random opening polymerization rate affects the ring opening dynamics(see Fig. 5 (c) and 5(d)).

**Figure 5.**
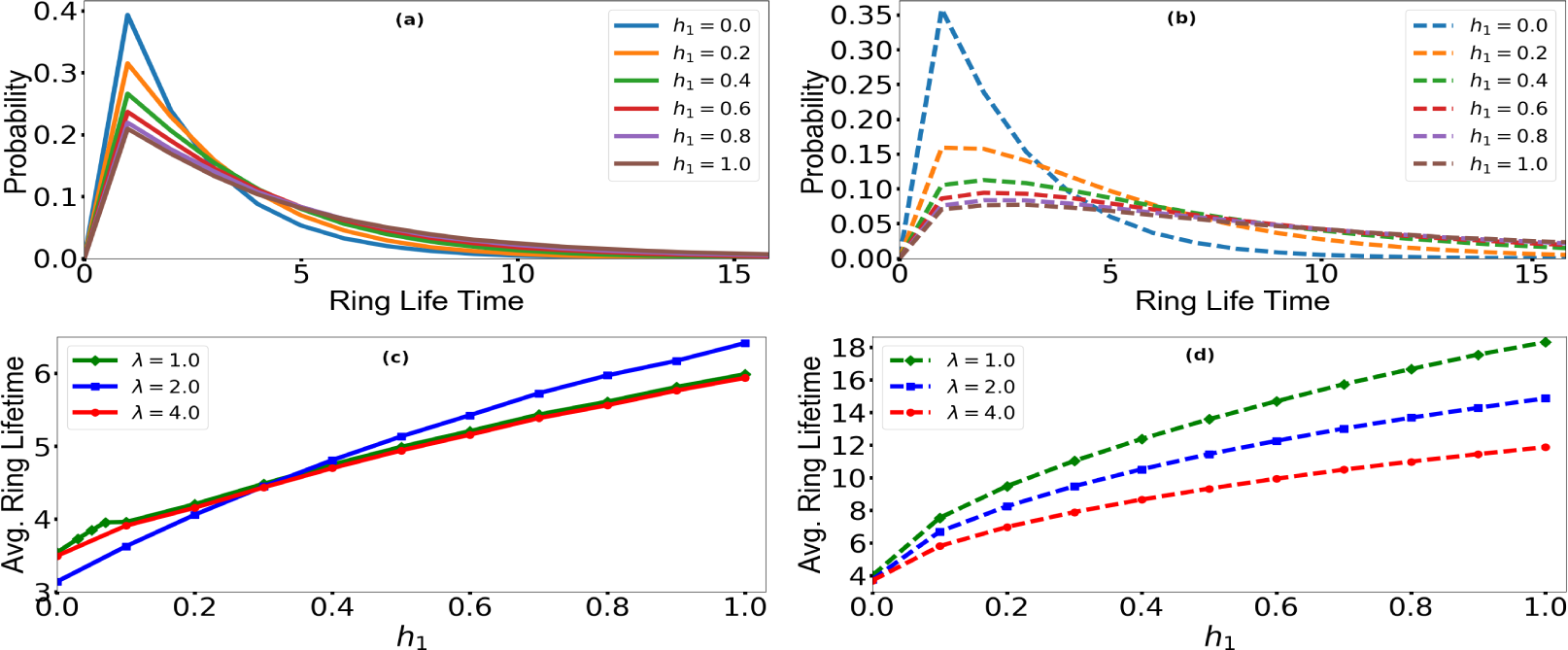
Distribution of ring length(top) in ring form for polymerization rate 4.0 and average length (bottom) for different polymerization rates for random (a and c) and nonrandom (b and d) breaking.

#### 3.1.4 Phase diagram for random breaking

From our observations on the average time for irreversible opening, it is clear that most stable ring is formed by fine tuning the hydrolysis rate in the case of vectorial hydrolysis with random breaking. We call this value of hydrolysis rate *h*_*critical*_, and find that it changes with the polymerization rate(see Fig 6). *h_critica_l* is the value where the life time distribution has peak in the Fig 4.

**Figure 6.**
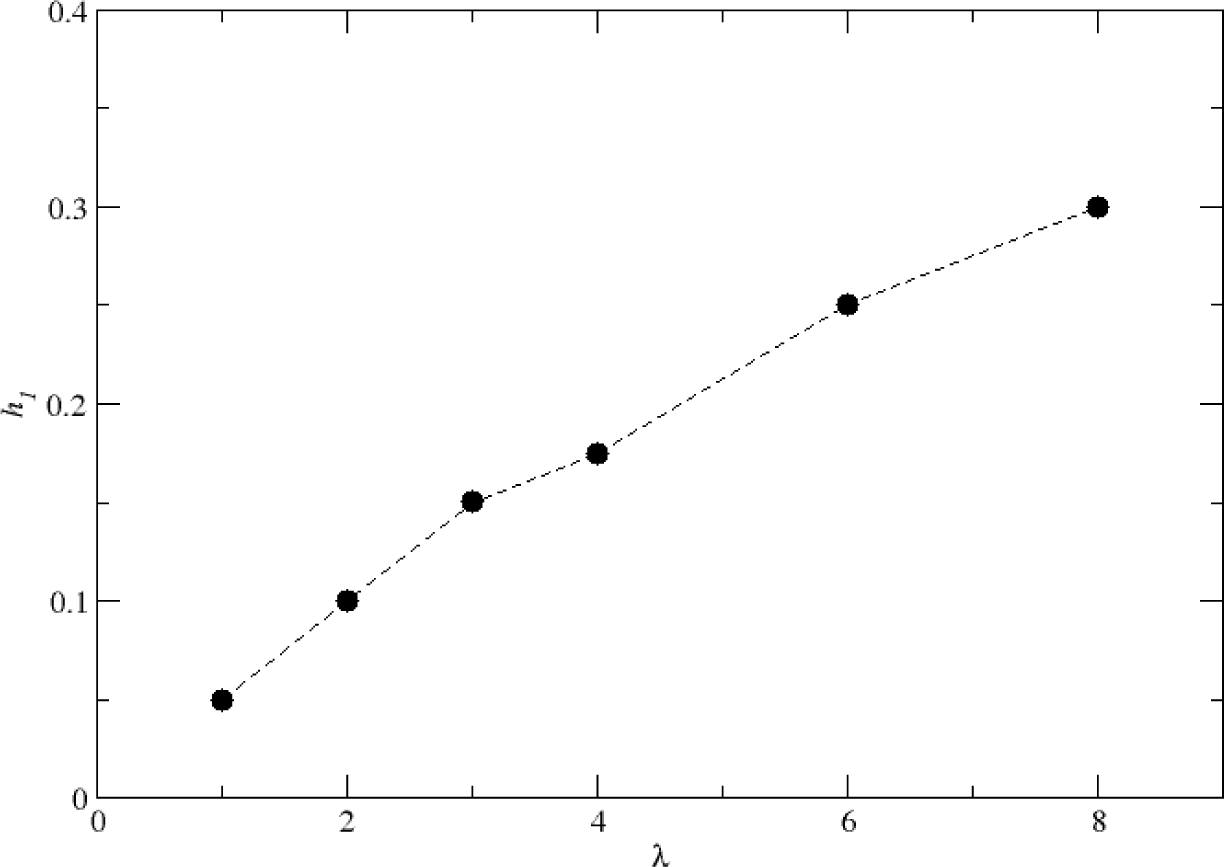
Phase diagram for random ring opening

### 3.2 Force generation and the effect of ring breaking rate on the dynamics

Based on the role played by contractile ring formed by filamentous actin and myosin motor proteins in creating the furrow that divides the eukaryotic cells into two during cytokinesis [56], it was proposed that the Z-ring plays a similar role in prokaryotic cells with conformation change in FtsZ due to hydrolysis creating the membrane constricting force[57]. The observation that curved FtsZ filaments create bending force on membranes[58] led to the hypothesis that bending of FtsZ protofilaments creates the force necessary for cell wall invagination and constriction in cells. A number of models were proposed to explain force generation in FtsZ [59, 60]. Though experiments could confirm that FtsZ filaments indeed generated force which could bend membranes and also showed vortex dynamics[25], the question remained that if this force was sufficient to constrict the membrane and the cell wall. More recent studies have shown that FtsZ treadmilling regulates cell wall formation[8, 9]. There are multiple independent units which synthesize Peptidoglycan(PD) and these units move along with the treadmilling FtsZ. FtsZ filaments deform the membrane and the PD synthesizing units add cell wall to the deformed region reinforcing it. This process leads to concentric rings of cell wall being formed and the radius of these rings constantly keeps on decreasing[9]. The next question is what would be the effect on the filament because of the PD synthesis mechanism it carries around and the local deformations of the membrane it is responsible for. In microtubules the force generation can be explained by pushing by the fibers[61]. But in the case of FtsZ the force is generated by the tension created in the filaments because of deformation in the membrane. This is due to the attached tethers and the movement of the synthesis machinery. The added stress can be thought as the increased probability of ring breaking. Hence one way to get the understanding of the force generation is by looking at the effect of ring breaking rate on the FtsZ dynamics.

Hence we studied the effect of ring breaking rate on the FtsZ ring for random breaking model with vectorial hydrolysis. As the breaking rates increased, the average length of the ring for a given hydrolysis and polymerization rate was higher(see Fig. 7). Also as the breaking rates increase, the average length for single ring breaking of the polymer for a given hydrolysis and polymerization rate was found to be lower. For polymerization rate 1.0 different breaking rates i.e 0.008 and 0.012 were found to be same. This suggests that for a given polymerization rate beyond a certain breaking rate, breaking rate does not have much effect on the average times for single ring breaking(see Fig. 8). We also looked at the average time of irreversible opening. We found that as the breaking rates increased, the hydrolysis rate at which the average time for irreversible opening becomes maximum was found to increase(Fig. 9). These plots qualitatively suggest that FtsZ ring with random breaking with vectorial hydrolysis is contractile in nature and is a viable model for force generation.

**Figure 7.**
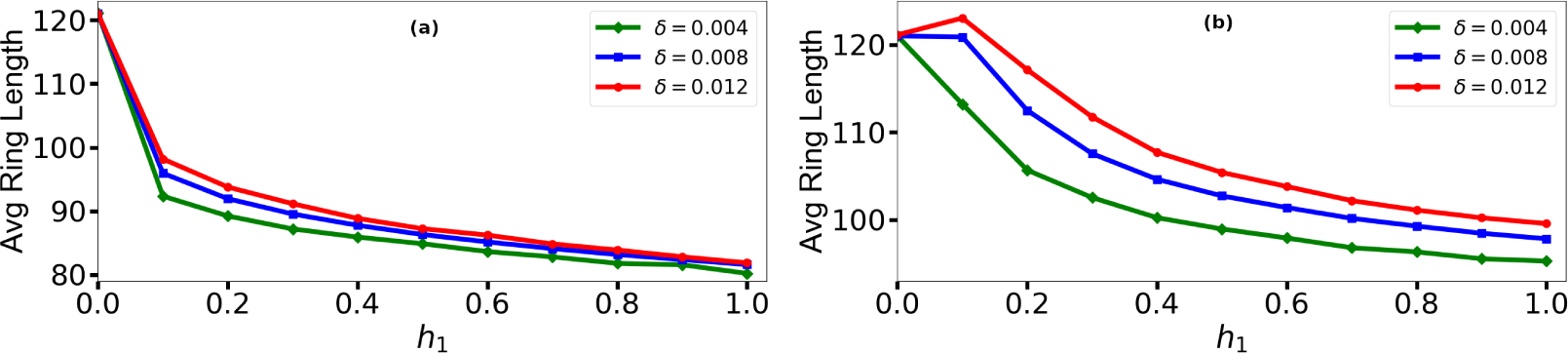
Average lengths for different ring breaking rates for different values of hydrolysis rates for polymerization rates 1.0(a) and 4.0 (b) for random breaking

**Figure 8.**
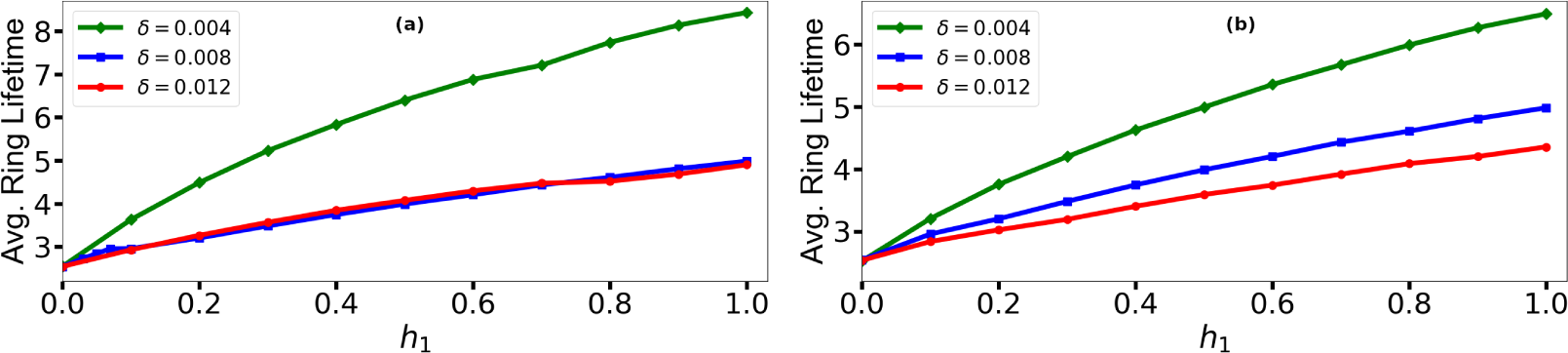
Average ring-life time for single break for different ring breaking rates for different values of hydrolysis rates for polymerization rates 1.0 (a) and 4.0 (b) for random breaking

**Figure 9.**
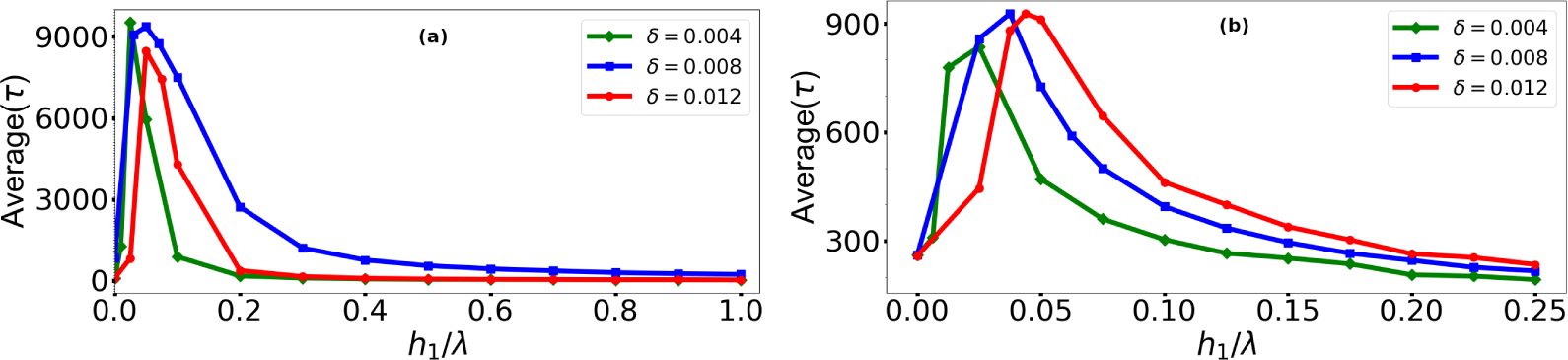
Average lifetime for irreversible opening for polymerisation rates 1.0 (a) and 4.0 (b).

### 3.3 Stochastic hydrolysis(SH)

SH in the case of microtubules provides a build-in mechanism for rescue and catastrophe and hence for the dynamics. This feature makes SH more likely than VH for microtubules. In this section we look at the FtsZ ring with SH. We will hence put *h*_1_ = 0 now.

We have seen in the previous section, the ring breaking naturally provides a way to build rescue and catastrophe in the FtsZ ring. For the sake of completion, we explore FtsZ dynamics assuming SH in this section. Fig 10 shows trajectory with both random breaking and non-random breaking. In the case of random breaking, there are now two ways to introduce interfaces in the system. As expected, the ring is more dynamic in this case and at the critical value of the hydrolysis rate *h*_2_, the ring stays stable for a very long time with regular opening and closing of the ring. In the case of non-random breaking the ring is similar, but the ring is not as stable as in the case of SH with random breaking.

We looked at the ring length distribution in this case. We find that the ring length distribution is sensitive to change of hydrolysis rate for both, random and non-random breaking(see Fig. 11). Though the non-random breaking case is less sensitive to the hydrolysis rate(especially for smaller polymerization rate).

We also looked at the time for irreversible ring opening and interestingly we find that unlike the case of vectorial hydrolysis, the average lifetime is insensitive to hydrolysis rate for a wide range of hydrolysis rate values.

**Figure 10.**
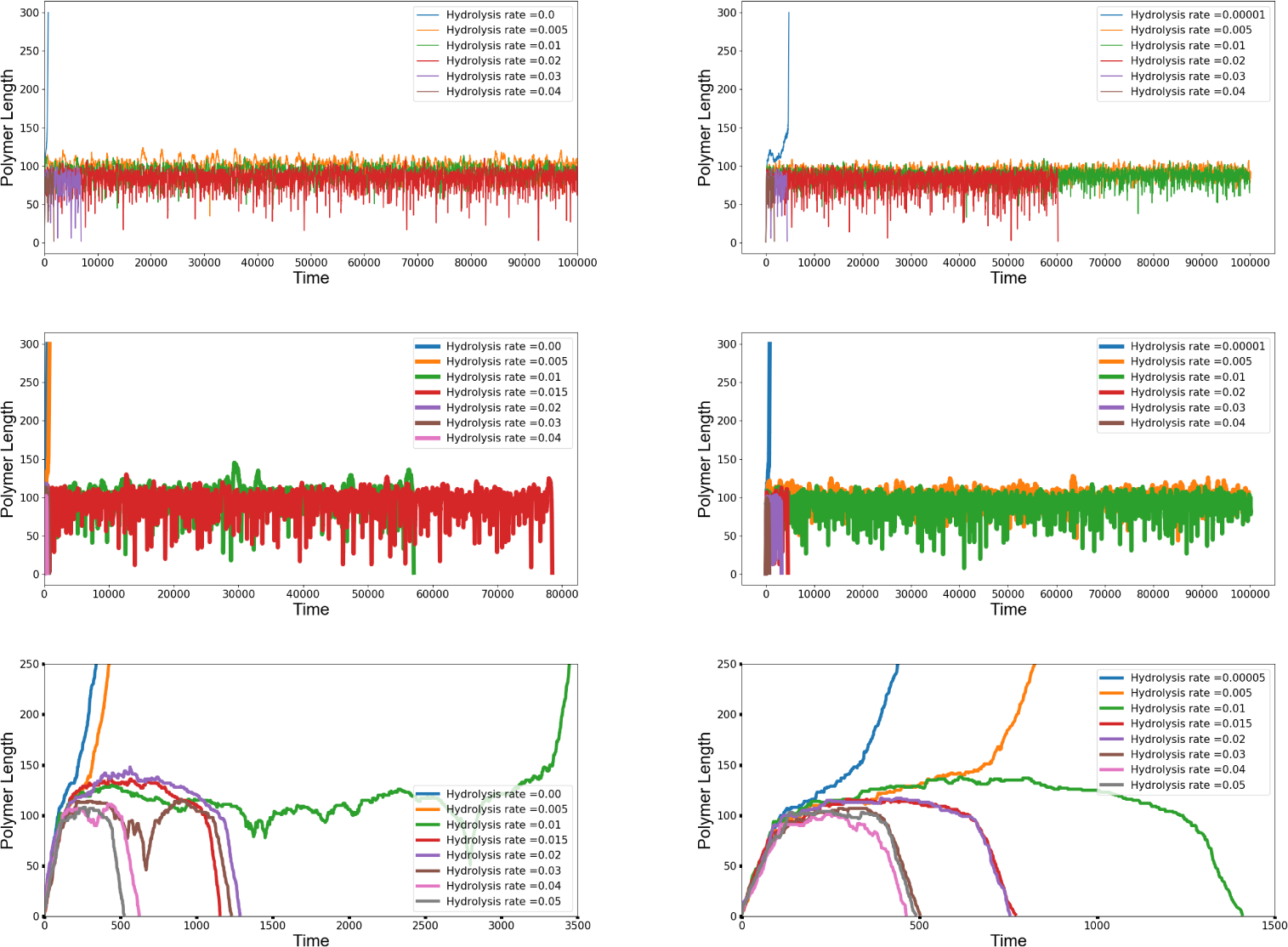
Trajectories for random (left) and non-random breaking (right) for polymerization rate 2.0, 4.0 and 8.0 respectively (top to bottom)

**Figure 11.**
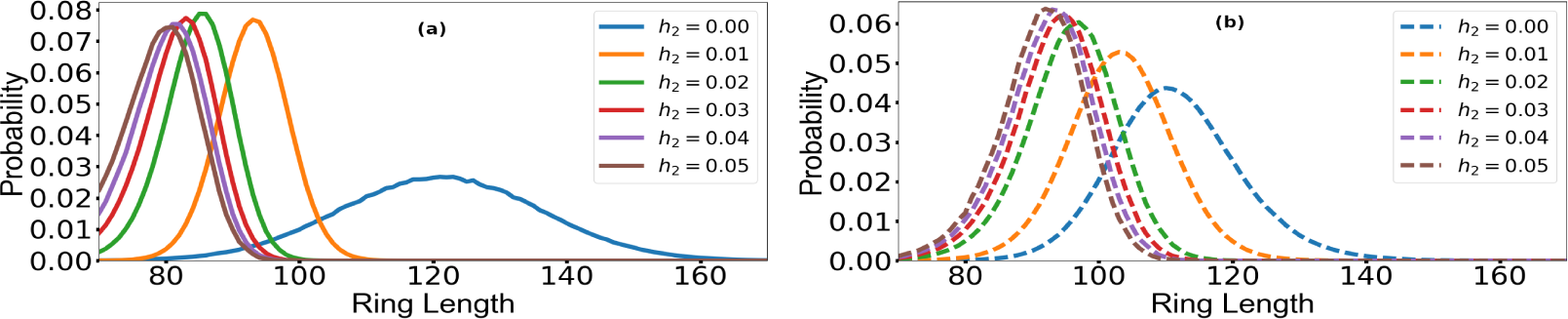
Distribution of ring length in ring form for polymerization rate 4.0 for random(a) and non-random (b) breaking.

**Figure 12.**
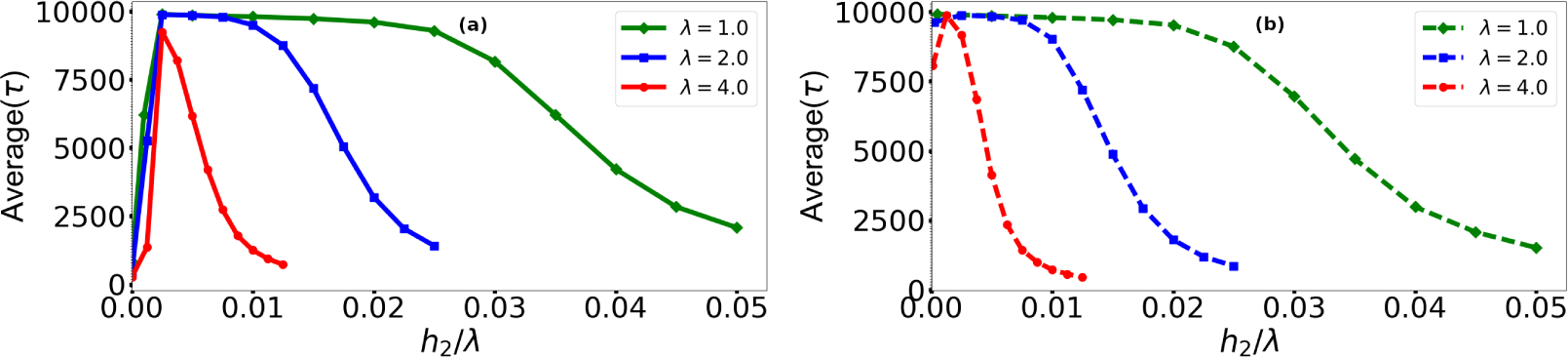
Average lifetime for irreversible opening for random (a) and non-random (b) breaking

## 4 Conclusion

The dynamics of FtsZ ring is being very actively studied in experiments in recent times[6–9]. It is still not possible to observe the microscopic dynamics and one can at best study the average quantities like the average lifetime and length of the ring [12]. In this paper, we have introduced a stochastic model which considers all the essential processes and compares all possible scenarios. This is important as the nature of hydrolysis and ring breaking is still not understood, even though there is a general consensus that hydrolysis and treadmilling of the FtsZ polymer plays a crucial role of defining its dynamics and functionality. Our analysis gives clear differences in the trend of measurable quantities. Note that the dynamic nature of the ring cannot be captured effectively in a deterministic model. Our stochastic model naturally gives rise to catastrophe and rescue of FtsZ ring, which have been observed in experiments. We find that unlike microtubules and actin the nature of hydrolysis does not play a crucial role for FtsZ ring. For microtubules and actin if one considers vectorial hydrolysis, then one has to introduce rescues by hand for the polymer to be dynamic. We find that the rescues and catastrophe events are built in our model due to the ring topology irrespective of the hydrolysis mechanism, suggesting ring breaking as a natural mechanism for introducing dynamics in the ring.

In the case of vectorial hydrolysis with random ring breaking we could qualitatively reproduce the known behavior of the *Z*-ring. We found that increasing the hydrolysis rate in this case narrows the ring length distribution and the average length of the ring decreases with hydrolysis. This is consistent with the invitro studies of Z-ring in [12]. We find that the time lapsed between ring formation and final decay(time for irreversible ring opening) changes non-trivially with the hydrolysis rate. One needs to fine tune hydrolysis rate to achieve the most stable ring in this case. In contrast, when we consider non-random ring breaking, the behavior of the ring is insensitive to the hydrolysis rate. Initially the polymer has only one interface where the hydrolysis occurs. But after ring closure, we find that in the case of random ring breaking, many new interfaces show up, and hence the ring can potentially break up at multiple places, consistent with the recent experiments [9]. In contrast in the case of non-random ring breaking, the number of interfaces do not change with time and one typically has only one interface in the *Z*-ring. All these observations suggests that *Z*-ring breaks randomly. Similar conclusion was also reached by [12]. Recent works suggest that treadmilling of *Z*-ring is sufficient to generate the constriction force generated by *Z*-ring during cell division. The ring with random breaking is contractile as shown in Section 3.2.

We have tried to keep our model basic so that we could distinguish the effect of different random processes clearly. The model can be studied by including many other processes like allowing the attachment/detachment of short polymer and not just a single monomer during polymerization/ depolymerization process. We find that this does not change the picture qualitatively and hence not needed to understand the basic dynamics of FtsZ polymer. For the sake of completion we also looked at the ring dynamics with stochastic hydrolysis. In this case we find that for both ring breaking mechanisms (random and non-random), we do not need to fine tune the hydrolysis rate to acheive the stable dynamic ring. Recent experiments [11] suggest that fine tuning of hydrolysis rate is required for stable dynamic ring. Behaviour of ring life time as a function of hydrolysis rate can hence be a distinguishing feature between vectorial and stochastic hydrolysis. Hence, differences in the statistics of the ring can be used to unravel the actual mechanism of hydrolysis in the FtsZ polymer.

We find that the randomness makes the ring more dynamic and stable, as seen in the case of stochastic hydrolysis and random breaking. In general it is now established that the stochasticity increases the stability of biological processes [62] and is important to understand the functioning of biological systems. Though the Z-ring has been studied by others using deterministic models, this is the first stochastic model for Z-ring We find that the model is able to explain a number of features of FtsZ dynamics, elucidating the crucial role the ring topology plays in the functioning of the *Z*-ring. The model successfully brings out the role of hydrolysis and ring breaking mechanisms in forming a contractile dynamic ring. The model is simple and in future can easily be used for more quantitative studies.

## Acknowledgements

The authors thank Dr. R. Sreenivasan and Dr. S. Roychowdhury for fruitful discussions. This work has been supported by Department of Atomic Energy, India thorugh the 12th plan project (12-R&D-NIS-5.02-0100).

## Author Contributions

A.S ran the simulations. S and A.V.A.K developed the model. A.S,S and A.V.A.K analysed the results and drafted the manuscript.

